# CENP-A drives asymmetric cell division and maintains stem identity

**DOI:** 10.1101/631598

**Authors:** Anna A. Dattoli, Ben L. Carty, Antje M. Kochendoerfer, Annie E. Walshe, Elaine M. Dunleavy

## Abstract

Centromeres, chromosomal loci essential for genome integrity, are epigenetically defined by CENP-A-containing chromatin. Recent studies suggest that parental CENP-A is asymmetrically distributed upon stem cell asymmetric division. However, a direct link between centromeres and stem cell identity has not been demonstrated. We show that *Drosophila* female germline stem cells (GSCs) and neuroblasts assemble centromeres between G2-phase and prophase, requiring CYCLIN A. Intriguingly, chromosomes that will be inherited by GSCs incorporate more CENP-A and capture more spindle fibers at pro-metaphase. Furthermore, over-expression of CAL1 (*Drosophila* CENP-A assembly factor) causes GSC-like tumours, while over-expression of both CENP-A and CAL1 promotes stem cell self-renewal. Finally, once centromeres have been assembled in GSCs, continued CENP-A assembly is not required in differentiating cells outside of the niche and CAL1 becomes dispensable. According to our results CENP-A regulates stem cell identity/maintenance. Moreover, crucial centromere assembly occurs in the niche prior to oocyte meiosis.

## INTRODUCTION

Stem cells are fundamental for the generation of all tissues during embryogenesis and replace lost or damaged cells throughout the life of an organism. During division, stem cells generate two cells with distinct fates: 1) a cell that is an exact copy of its precursor (maintaining the “stemness”) and 2) a daughter cell that will subsequently differentiate (Betschinger and Knoblich, 2004; Inaba and Yamashita, 2012). Epigenetic mechanisms (heritable chemical modifications of the DNA/nucleosome that do not alter the primary genomic nucleotide sequence) were recently discovered to regulate the process of self-renewal and differentiation of stem cells (Christophersen and Helin, 2010; Eun et al., 2010). In *Drosophila* male GSCs phosphorylation at threonine 3 of histone H3 (H3T3P) preferentially associates with chromosomes that are inherited by the future stem cell prior to segregation (Xie et al., 2015a). Furthermore, centromeric proteins seem to be asymmetrically distributed between stem and daughter cells upon division in the *Drosophila* intestine (García del Arco et al., 2018). These findings support the silent sister hypothesis (Lansdorp, 2007), according to which epigenetic variations differentially mark sister chromatids driving selective chromosome segregation during stem cell mitosis (Caperta et al., 2008; Dai et al., 2005; Lansdorp, 2007; Tran et al., 2013; Xie et al., 2015a). Centromeres, the primary constriction of chromosomes, are crucial for genomic integrity providing the chromatin surface where the kinetochore assembles (McKinley and Cheeseman, 2016). In turn, the kinetochore ensures the correct attachment of spindle microtubules and faithful chromosome partition into the two daughter cells upon division (Musacchio and Desai, 2017). Centromeric chromatin contains different kind of DNA repeats (satellite and centromeric retrotransposons) (Fukagawa and Earnshaw, 2014) combined with nucleosomes containing the histone H3 variant CENP-A. A common feature of centromeres is that they do not possess any “password”, in terms of DNA sequence, involved in their specification. Rather, they are specified epigenetically by CENP-A (Allshire and Karpen, 2008; Black and Cleveland, 2011; Fukagawa and Earnshaw, 2014). Centromere assembly, classically measured as CENP-A deposition to generate the centromeric nucleosomes occurs at the end of mitosis (between telophase and G1) in humans (Hemmerich et al., 2008; Jansen et al., 2007). Interestingly, additional cell cycle timings for centromere assembly have been reported in flies (Ahmad and Henikoff, 2001; Mellone et al., 2011; Schuh et al., 2007). Specifically, *Drosophila* spermatocytes and starfish oocytes are the only cells to date to assemble centromeres prior to chromosome segregation, during prophase of meiosis I in an animal tissue (Dunleavy et al., 2012; Raychaudhuri et al., 2012; Swartz et al., 2018). These examples show that centromere assembly dynamics can differ among metazoans and also among different cell types in the same organism.

A key player in centromere assembly in vertebrates is HJURP (Holliday Junction Recognition Protein), which localises at the centromeres during the cell cycle at the same time window proposed for CENP-A deposition (Dunleavy et al., 2009; Foltz et al., 2009). Furthermore, centromere assembly is due to the interplay between cell cycle regulators and the assembly machinery. In flies, CID (the homologue of CENP-A) deposition requires activation of the anaphase promoting complex/cyclosome (APC/C) and degradation of CYCA (Mellone et al., 2011), while centromere assembly is antagonised by Cdk1 activity and promoted by the kinase Plk1 in humans (McKinley and Cheeseman, 2014; Silva et al., 2012; Stankovic et al., 2017a). To date, little is known about centromere assembly dynamics and its possible role during stem cell asymmetric division. Indeed, *Drosophila melanogaster* ovaries provide an excellent model to study centromere dynamics in stem cells in their native niche (Yan et al., 2014). In this tissue germ stem cells (GSCs) are easily accessible and can be manipulated genetically. Moreover, the cell cycle assembly time of centromeres in GSCs and their differentiated cells, cystoblasts (CBs), could be used as a mechanism to epigenetically discriminate between these two cell types. In *Drosophila*, CID binds to CAL1 (fly functional homologue of HJURP) in a prenucleosomal complex and its localisation to the centromeres requires CAL1 and CENP-C (Erhardt et al., 2008; Mellone et al., 2011).

Taking advantage of this model system together with confocal and super-resolution microscopy, we investigated the dynamics of centromere assembly throughout the cell cycle in GSCs and CBs. We show that centromeres are assembled between G2 and prophase in both GSCs and CBs. Furthermore, neuroblasts follow the same trend. Moreover, CYCA localises at the centromeres in GSCs and its knock down is responsible for a marked reduction of centromeric CID and CENP-C, but not CAL1. Our super-resolution analysis of GSCs at prometaphase shows that chromosomes that will be inherited by the stem cell are loaded with more CID. This asymmetry is not observed in differentiated cells outside of the niche. Intriguingly, GSC-chromosomes anchored more spindle fibers. Furthermore, while CID knock down results in agametic ovaries, depletion of CAL1 at centromeres blocks cell GSC proliferation and differentiation. Surprisingly, over-expression of CID and CAL1 together promotes stem cell self-renewal, while over-expression of CAL1 alone causes GSC-like tumours. CAL1 and CID knock down at later stages of egg development have no obvious effect on cell division because these cells inherit CID from GSCs. Taken together, our findings suggest a strong link between CID and stem cell identity, while CAL1 drives cell proliferation.

## RESULTS

### Nuclear distribution of centromeres in GSCs changes through the cell cycle

The cell cycle assembly time of centromeres in GSCs and CBs is currently unknown. To elucidate this process, we observed the distribution of centromeres throughout the cell cycle. The *Drosophila* GSC niche is found at the apical end of the germarium, the anterior tip of the adult ovariole (Fig. 1A, region 1). The niche comprises the terminal filament and usually 2-3 GSCs attached to the cap cells via a cytoplasmic structure, the spectrosome. The spectrosome is a bridge, which allows BMP (Bone Morphogenetic Protein) signalling to be transferred to the stem cells and induce cell division (Mottier-Pavie et al., 2016). Furthermore, its shape can be used to define the cell cycle stage of the GSC (Ables and Drummond-Barbosa, 2013; Kao et al., 2015). Upon asymmetric division, the new daughter cell closer to the niche retains the “stemness”, while the other, the CB differentiates. GSCs and CBs are easily distinguishable by distinct morphological features and anatomical position in region 1 (Figure 1A). Each CB undergoes four rounds of mitosis with incomplete cytokinesis, giving rise to 16-cell cysts of cystocytes (CCs) interconnected to each other through the fusome, a branched spectrosome. After completion of pre-meiotic S phase, all of the 16 cysts start meiosis and form the synaptonemal complex. The oocyte originates from either of the two CCs with four fusome-bridges (Fig. 1A, region 2a-2b, brown cells) (Christophorou et al., 2013; Rangan et al., 2011). In region 3, the 16-cell cysts will mature to an egg chamber containing 15 nurse cells that provide for the oocyte (Fig. 1A, region 3), which completes meiosis (Hughes et al., 2018; McLaughlin and Bratu, 2015).

**Figure 1.**
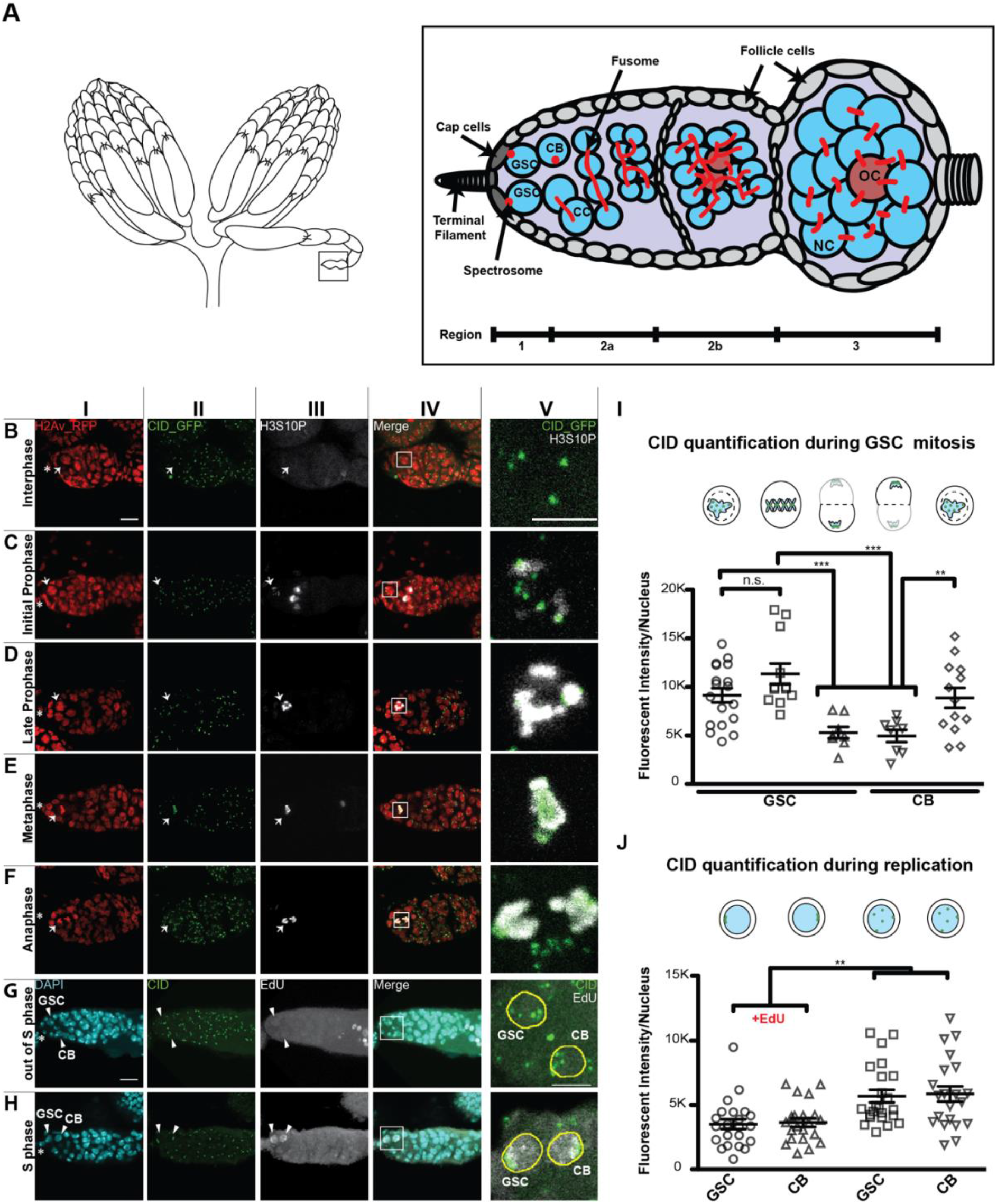
Centromere assembly occurs between G2 and prophase in *Drosophila* female GSCs. **A**) Diagram of *Drosophila* ovary (left) and germarium containing the germline stem cell niche (right). GSC= germline stem cell; CB= cystoblast; CC= cystocyte; NC= nurse cells; OC= oocyte. The spectrosome (red) connects GSCs to the cap cells (dark grey). **B-F**) Confocal z-stack projection of a germarium expressing H2AV-RFP (red) (**I**), GFP-CID (green) (**II**) and stained for H3S10P (white) (**III**) showing centromere localisation in GSC nuclei throughout the cell cycle. (**V**) shows inset marked by box in merged image (**IV**). **B**) Interphase **C-D**) Initial and late prophase **E**) Metaphase **F**) Anaphase **G**) EdU negative (out of S phase) GSCs and CBs. **H**) EdU positive (S phase, white) GSC and CB. **I-J**) Star indicates the position of the terminal filament; arrows indicate GSC position in the germarium; arrow heads indicate the position of GSC and CB in the germarium; **B-F**) < 1 day old female flies heterozygous transgenic flies CID-GFP/H2Av-RFP. **G-H**) 1-3 days old wild type female flies; **I-IV** scale bar 10 μm; **V** scale bar 5 μm. Quantification of the fluorescence intensity of CID observed at centromeres, obtained through the expression of CID-GFP (**I**) or antibody staining (**J**). Fluorescence Intensity is expressed as mean grey values after background subtraction (see material and methods); data are represented as the mean±standard error of the mean (SEM); **= p value<0.005; ***= p value<0.0005.

To our aim, we used transgenic flies expressing CID coupled to GFP, to follow centromeres, and H2Av coupled to RFP (Schuh et al., 2007) to follow chromatin condensation. To identify each phase of mitosis in GSCs we used the phosphorylation at serine 10 of histone H3 (H3S10P) (Hendzel et al., 1997; Matias et al., 2015). At interphase chromatin is not condensed (Fig. 1BI), centromeres are spread throughout the nucleus (Fig. 1BII, 1BV) and H3S10P signal is absent (Fig. 1BIII, 3BV). At prophase, H3S10P signal is present, chromosomes begin to condense and centromeres start to align on the metaphase plate (Fig. 1C-D). At this stage we observed on average 5.7 centromere foci per cell. At metaphase, chromosomes and centromeres are completely aligned on the metaphase plate (Fig. 1EI and EII). At this stage it is possible to clearly distinguish the centromeres of each set of sister chromatids that will be inherited respectively by the new GSC and CB (Fig. 1EII, EV). We observed an average of 6.9 centromere foci aligned on the metaphase plate. At anaphase, chromosomes and centromeres migrate to the opposite pole of each new daughter cell (Fig. 1FI, FII and FV). At this point an average of 3.1 centromere foci are visible per cell. At telophase, the H3S10P signal is notably reduced, the chromatin starts to de-condense and centromeres are still located at the opposite side of each nucleus of the new GSC and CB (not shown).

To identify cells which were in or out of S phase, we used 5-ethynyl-2′-deoxyuridine (EdU) to label nuclei with or without newly replicated DNA (Salic and Mitchison, 2008). After EdU incorporation and fixation, ovaries were antibody-stained to study centromere positioning in GSCs and CBs during replication (Fig. 1G-H). In EdU negative cells, DNA is not condensed and centromeres are scattered throughout the nucleus (Fig. 1GI-V), similar to the interphase pattern (Fig.1BI-V). These cells show on average 4.6 centromere foci per nucleus. Interestingly, approximately 91% of the cells analysed (40/44) show that GSCs and CBs were simultaneously positive for the EdU staining with approximately 3.4 centromere foci (Fig. 1HI-V). In these cells, centromeres assumed a similar localisation to that observed during anaphase and telophase of GSC division, localising to the opposite poles of the GSC and CB nuclei (Fig. 1FV and 1HV). With the aid of several cell cycle markers (FUCCI) we did not succeed to isolate the G1 stage (data not shown), suggesting that it is very short in GSCs as previously proposed (Ables and Drummond-Barbosa, 2013). This together with the DNA condensation and centromere distribution, allow us to conclude that cells that were EdU negative are likely to be between G2 phase and initial prophase of the cell cycle. In summary, our cell cycle analysis of centromere localisation in M and S phases shows that centromeres are localised at the opposite poles of the new GSC and CB nuclei at anaphase, and that during DNA replication centromeres retain this localisation.

### Centromeric recruitment of CID occurs between G2 and prophase in both *Drosophila* GSCs and neuroblasts

To assess the cell cycle timing of centromere assembly in GSCs, we quantified the CID fluorescent intensity in mitosis and interphase. We first quantified the total amount of CID-GFP per nucleus at each phase of mitosis using the H3S10P marker (Fig. 1I). No significant difference between CID levels was detected at prophase (GSC_p_=9142±726.9, n=18 cells); and metaphase (GSC_m_=11334±1073, n=12 cells). Furthermore, the values at prophase for GSCs are not dissimilar from the CB (CB_p_=8882±1024, n=13 cells). As expected, compared to metaphase, during anaphase CID levels drop about half with comparable fluorescent intensity values in both daughter cells (GSC_a_=5302±583, n=8 cells and CB_a_=4965±673, n=8 cells). We then quantified the amount of CID, through antibody staining, in EdU positive (S phase) or negative (G2 phase/prophase) cells (Fig.1J). Surprisingly, the total amount of CID detected per nucleus in S phase cells was significantly lower than the one obtained for G2 phase/prophase stage in both GSC and CB, exhibiting the following values of fluorescent intensity: GSC_+EdU_=3502±386 (n=23 cells), CB_+EdU_=3642±345 (n=21 cells); GSC_-EdU_=5690±486 (n=23 cells), CB_−EdU_=5863±597 (n=21 cells). These results show that low levels of CID are observed during anaphase and replication, while considerably higher levels of CID are measured during G2/prophase, prophase and metaphase, suggesting that centromere assembly likely initiates during G2 phase in *Drosophila* female GSCs. Furthermore, gradual deposition of CID continues up to prophase and remains stable at metaphase. Finally, a similar trend was observed for the CBs.

To exclude the possibility that the centromere assembly in G2 phase was a specific feature of GSCs only, we investigated CID deposition in reactivated neuroblasts (NBs) of the thoracic ventral nerve chord (tVNC, Fig. 2A) in larval brains. To isolate neural stem cells in G2/prophase, we antibody-stained with the NB marker Deadpan (DPN) (Boone and Doe, 2008) and the G2 regulator CYCLIN A (CYCA) (Fig. 2K) that is crucial for the G2-M transition and is degraded at metaphase (Lilly et al., 2000). DPN positive NBs display different sizes, between 4 and 8 μm (Fig. 2B), as already described in literature (Chell and Brand, 2010). We then quantified the total amount of CID per nucleus in these cells, through antibody staining (Fig. 2C). In the CYCA negative NBs, the DNA is not condensed, indicating that they are neither in mitosis nor in G2/prophase. We therefore labelled them as G1/S-phase NBs. Our quantification shows that CYCA positive NBs have approximately 40% more CID compared to the G1/S-phase NBs (G2/prophase=4190±364, n=30 cells; G1/S-phase (4 μm)=2191±151, n=31 cells; G1/S-phase (5-8 μm)=2552±155, n=30 cells; 9 tVNC analysed). Our results confirm that similar to GSCs also neural stem cells assemble centromeres between G2 and prophase.

**Figure 2.**
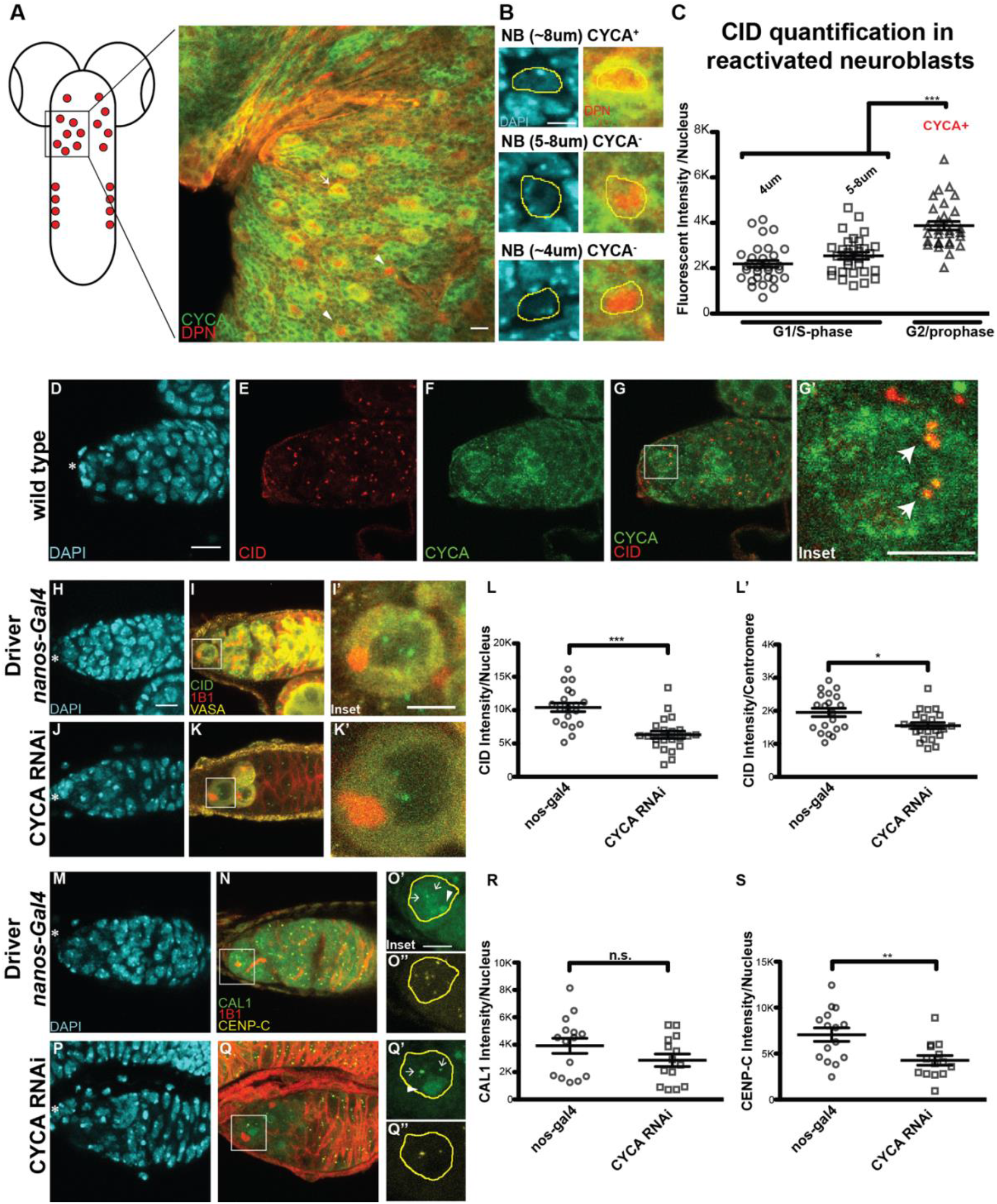
CID and CENP-C deposition requires CYCLIN A in *Drosophila* female GSCs. **A**) Diagram of *Drosophila* larval brain containing neuroblasts (red) and confocal z-stack projection of a section of the thoracic ventral nerve chord (tVNC) stained with for DAPI (cyan), anti-CYCA (green), anti-DPN (red) and anti-CID (not shown). **B**) Neuroblast (NB) in the tVNC are present in different sizes. **C**) Quantification of fluorescence intensity of CID at centromeres in CYCA negative and positive NBs. Fluorescence Intensity is expressed as integrated density after background subtraction (see material and methods). **D-G’**) Germarium of wild type flies stained for DAPI (cyan), anti-CID (red) and anti-CYCA (green). **H-K’**) Confocal z-stack projection of a germarium of *nanos-Gal4* (**H-I’**) and CYCA RNAi flies (**J-K’**), stained for DAPI (cyan), anti-VASA (yellow), anti-CID (green) and anti-1B1 (spectrosome, red). **L-L’**) Quantification of fluorescence intensity of CID observed at centromeres per nucleus (**L**) and per individual centromere averaged per each nucleus (**L’**). **M-Q**”) Confocal z-stack projection of a germarium of *nanos-Gal4* (**M-O**”) and CYCA RNAi flies (**P-Q**”), stained for DAPI (blue), anti-CAL1 (green), anti-CENP-C (yellow) and anti-1B1 (spectrosome, red). **R**) Quantification of fluorescence intensity of centromeric CAL1 per nucleus, using CENP-C as a centromeric marker based on two biological replicates. **S**) Quantification of fluorescence intensity of CENP-C per nucleus, based on two biological replicates. Fluorescence Intensity is expressed as mean grey values after background subtraction (see material and methods); data are represented as the mean±standard error of the mean (SEM); ***= p value<0.0005; *= p value< 0.05. Star indicates the position of the terminal filament, arrows indicate centromeres and arrow heads indicate the nucleolus; 3 days old female flies; scale bar 10 μm; inset scale bar 5 μm.

### CID loading at GSC centromeres requires CYCLIN A

Previous work showed that centromere assembly is tightly linked to key cell cycle regulators (Stankovic *et al*., 2017). In *Drosophila*, CYCA accumulation in G2 phase is crucial to make centromeres competent for assembly, while it needs to be degraded by the proteasome activity in order to allow CID deposition (Erhardt et al., 2008; Mellone et al., 2011). Our work shows that in GSCs, centromeric recruitment of CID initiates in G2 phase and continues until prophase (Fig. 1), coinciding with CYCA activity. Therefore, we characterised the localisation pattern of CYCA in GSCs with respect to centromeres. CYCA was previously shown to have both a cytoplasmic and nuclear localisation, specifically co-localising with CID at the centromeres in Kc167 cells (Erhardt et al., 2008). We confirm that this is the case also for GSCs using antibody staining (Fig. 2D-G and inset G’). Following, we used the GAL4:UAS system (Duffy, 2002) to deplete CYCA specifically in GSCs. Thus, a double stranded RNA interference line for *cyclin a* was combined with the germline-specific driver *nanos-Gal4*, strongly expressed in GSCs (Mathieu et al., 2013). To confirm the knock down, control *nanos-Gal4* and CYCA RNAi ovaries were dissected and antibody stained against CYCA (Fig. S1). To identify possible phenotypes with respect to centromere assembly, we used antibody staining against germline (VASA) and stem cell (1B1, spectrosome) specific markers (Yan et al., 2014), in combination with centromeric markers. VASA staining showed that in control *nanos-Gal4* (F0), germaria chambers are filled with germ cells (Fig. 2H-I’) while CYCA depletion leads to a loss of germ cells (Fig. 2J-K’). Furthermore, the few germ cells left appear to be as twice as big as the germ cells in the control (Fig 2H-K’, focus on inset 2I’ and K’). We next quantified the amount of CID detected in GSCs (Fig. 2 insets I’ and K’, graphs L-L’). For this we specifically considered GSCs with a round spectrosome, attached to the cap cells, and with decondensed DNA, which defines them to be between G2 and early prophase (see Fig. 2 inset I’ and K’). We first observed that *nanos-Gal4* GSCs contain an average of 5.4 centromere foci detected with CID antibody, while GSCs depleted for CYCA show only 4. We then quantified total centromeric CID levels per nucleus in both of the samples, finding that in GSCs from CYCA-knocked down germaria these levels are reduced by 40% compared to the control (Fig. 2L, *nanos-Gal4*=10411±642, n=20 cells; CYCA RNAi= 6303±538.5, n=22 cells). This reduction in CID intensity was detected at each individual centromere also (Fig. 2L’). According to our data, CYCA RNAi-GSCs retain approximately 80% of CID at centromeres compared to the control (Fig.2L’, *nanos-Gal4*=1949±127, n=20 cells; CYCA RNAi= 1548±93.45, n=22 cells). Taken together, our data show that CYCA depletion is specifically responsible for the 40% loss of centromeric CID per nucleus and a 20% loss of CID at individual centromeres.

### CAL1 levels at the centromeres are not affected by CYCLIN A deficiency

To test whether the loss of CID observed in CYCA deficient GSCs was due to the loss of CAL1, and/or CENP-C, we antibody stained control and knocked down germaria for both CAL1 and CENP-C (Fig. 2M-Q). CAL1 is not only detectable at centromeres but also in the nucleolus (Schittenhelm et al., 2010; Unhavaithaya and Orr-Weaver, 2013). In this case, we specifically quantified the centromeric CAL1 in GSCs at G2/prophase and found no significant difference in the amount of centromeric CAL1 detected per nucleus between the *nanos-Gal4*-GSCs and CYCA RNAi-GSCs (Fig. 2R, *nanos-Gal4*=3921±546.4, n=15 cells; CYCA RNAi= 2865±457.8, n=14 cells). In contrast, CENP-C levels are reduced, comparable to CID (fig. 2S, *nanos-Gal4*=7060±730.1, n=15 cells; CYCA RNAi= 4269±525.6, n=14 cells). These results suggest that the diminishment of CENP-C and CID observed in the GSCs of CYCA-knocked down germaria might be independent of CAL1.

### Super-resolution imaging reveals that chromosomes to be inherited by the GSCs are loaded with more CID and capture more spindle fibers

To explore the biological significance of centromere assembly in G2-prophase and to provide a possible link to asymmetric divisions occurring in stem cells, we investigated CID intensity at sister chromatids in GSCs prior to division. Specifically, we used super resolution microscopy to examine CID intensity at sister centromeres at pro-metaphase and metaphase. To capture GSCs in this specific time-window, we used the phosphorylation at threonine 3 of histone H3 (H3T3P) (Xie *et al*., 2015) marker and antibody staining against the spectrosome, which has a round shape during mitosis (Ables and Drummond-Barbosa, 2013). Importantly, super-resolution microscopy allowed us to resolve 8 individual sister chromatids pairs at these stages (16 centromere foci) (Fig. 3). Using the position and orientation of the spectrosome, we specifically identified centromeres that will be inherited by the GSCs and centromeres that will belong to the CBs (Fig. 3A-F, Fig. S2). Following, we measured the total amount of CID present on one set of chromosomes versus the other. For comparison, we conducted the same analysis on differentiated cells of 4-cell cysts called cystocytes (CC) that divide symmetrically (Fig. 3G-K). The ratio obtained between values shows that centromeres present on the GSC side incorporate approximately 20% more CID, compared to centromeres of the CB side (ratio GSC_side_/CB_side_=1.192±0.072, n=9 GSCs in pro-metaphase/metaphase; Fig. 3L-M). Importantly, this asymmetry in the distribution of CID is not observed in the chromosomes of the CCs at the same time-window (ratio CCA_side_/CCB_side_=1.016±0.027, n=9 CC in pro-metaphase/metaphase; Fig. 3L-M). Because it was already shown that chromosome having bigger centromeres can dock more spindle fibers (Drpic et al., 2018), we decided to check whether this would be the case also with GSC-chromosome which harbour more CID. We therefore antibody stained GSCs in pro-metaphase and metaphase against tubulin (Fig. 3N-S). We first observed that at pro-metaphase the spindle is already bioriented (Fig. 3N-P). Secondly, we observed that the spindle fibers nucleated from the daughter centrosome, inherited by the GSC (Salzmann et al., 2014), are more compared to those nucleated from the mother centrosome on the CB side. This is detectable both at pro-metaphase and metaphase (Fig. 3N-S). These results suggest that chromosomes are labelled with differential amount of CID upon centromere assembly, and specifically chromosomes inherited by the GSCs harbour more CID. Furthermore, GSC-chromosomes in this time-window capture more spindle fibers compared to the CB-chromosomes.

**Figure 3.**
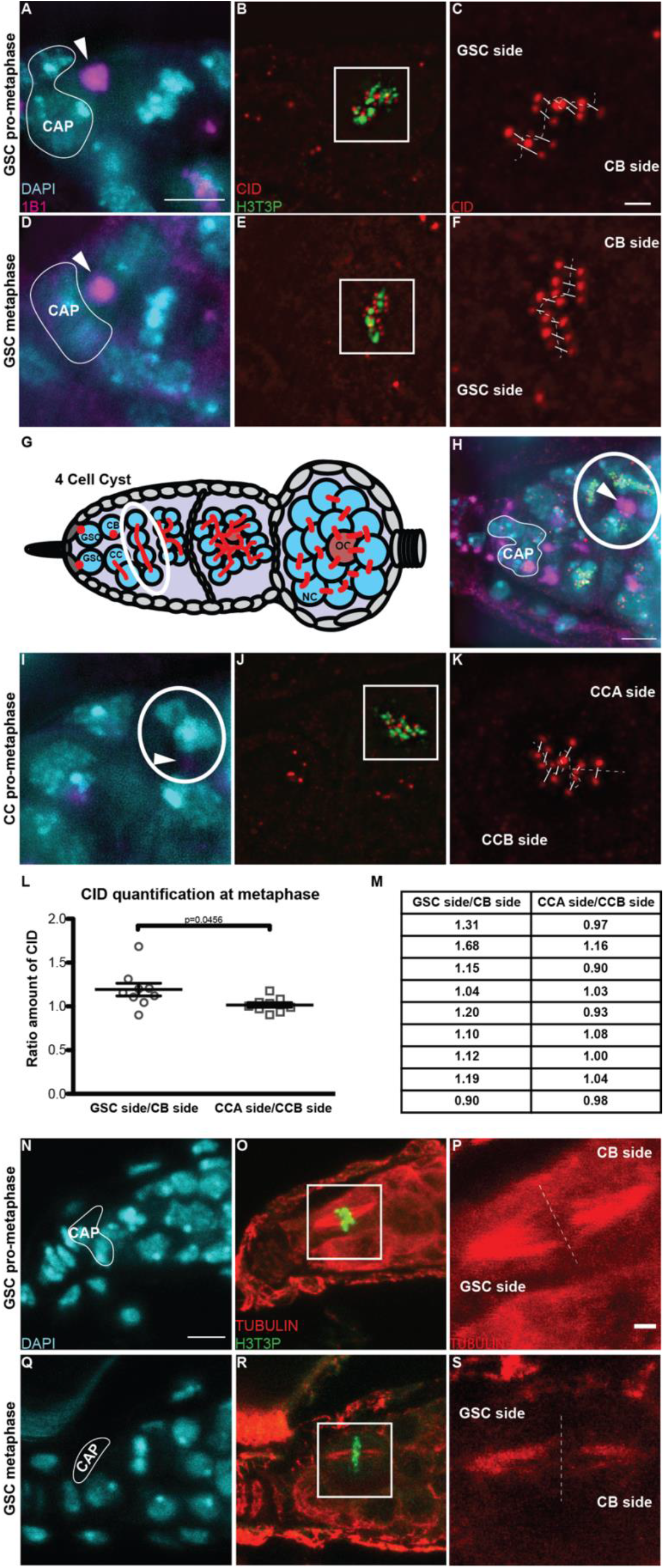
Chromosomes retain differential amount of CID upon centromere assembly in *Drosophila* female GSCs. **A-F**) Super-resolution (N-SIM) z-stack projection of a GSC at pro-metaphase (**A-C**) and metaphase (**D-F**). **G**) Diagram of *Drosophila* germarium, highlighting the 4-cell cyst stage containing 4 cystocytes (CCs). **H**) Super-resolution (N-SIM) z-stack projection of a germarium capturing 4 CCs in prometaphase/metaphase, which divide synchronously. **I-K**) Super-resolution (N-SIM) z-stack projection of a CC at prometaphase. **L**) Ratio between the total amount of CID detected on the chromosomes of the GSC side/ the total amount of CID detected on the chromosomes of the CB side, and similarly for CCA and CCB sides. **M**) Table of the ratio values obtained for each cell analysed. Ovaries were stained for DAPI (cyan), anti-1B1 (spectrosome, magenta), anti-CID (red) and anti-H3T3P (green). **N-S**) Confocal z-stack projection of a GSC at pro-metaphase (**N-P**) and metaphase (**Q-S**). Ovaries were stained for DAPI (cyan), anti-TUBULIN (red) and anti-H3T3P (green). Arrows indicate the spectrosome, arrow heads indicate the fusome (not yet visible in the z-stacks projected in **I-K**). 30 min old female flies; scale bar 5 μm; inset scale bar 1 μm.

### CAL1 is crucial for stem cell division and differentiation in *Drosophila* female germline tissues

To test the role of centromere assembly during stem cell asymmetric division, we performed functional analyses in GSCs. Specifically, we used UAS-RNAi for CID and CAL1 transgenic lines (Dietzl et al., 2007) combined with the driver *nanos-Gal4*. Ovaries from F1 progeny were dissected and screened using antibody staining against VASA and the spectrosome (1B1). Control ovaries showed germaria filled with germ cells (Fig. S3A) and stem cells having a round spectrosome attached to the cap cells (Fig. S3A, stylised arrows). As expected for an essential gene, CID knocked down female flies (F1) showed agametic ovaries with no VASA-positive cells (Fig. S3A) and these flies did not lay eggs (not shown). For the CAL1 knock down, we confirmed a greater than 10-fold depletion of CAL1 expression through real time-qPCR (Fig. 4A, see STAR methods). Phenotypic analysis of CAL1 depleted ovaries showed they were largely agametic (Fig. S3A). However, 18% of germaria (3/16) showed the presence of a few cells (1-3) that were VASA positive with a round spectrosome and located 90% of the time at the apical end germaria (Fig. S3A). Older flies (7 days post eclosion) showed a higher frequency of this phenotype (approximately 30%, 5/16 germaria) (not shown). The 1-3 cells left showed all the features of GSCs (Fig. 4BI-IV). They are VASA-positive (Fig. S3A), located at the apical end of the germarium close to the terminal filament in the stem cell niche, have a round spectrosome (Fig. S3A, compare 4BII and 4CII) and stain positive for phosphorylation of Mothers Against Dpp (pMAD), a BMP signaling indicator present in GSCs (Song, 2004) (compare Fig. 4BIII and 4CIII) and do not express the differentiation marker Bag of Marbles (BAM) (Fig. S3B). From our analysis we can conclude that CAL1 is crucial for stem cell differentiation.

**Figure 4.**
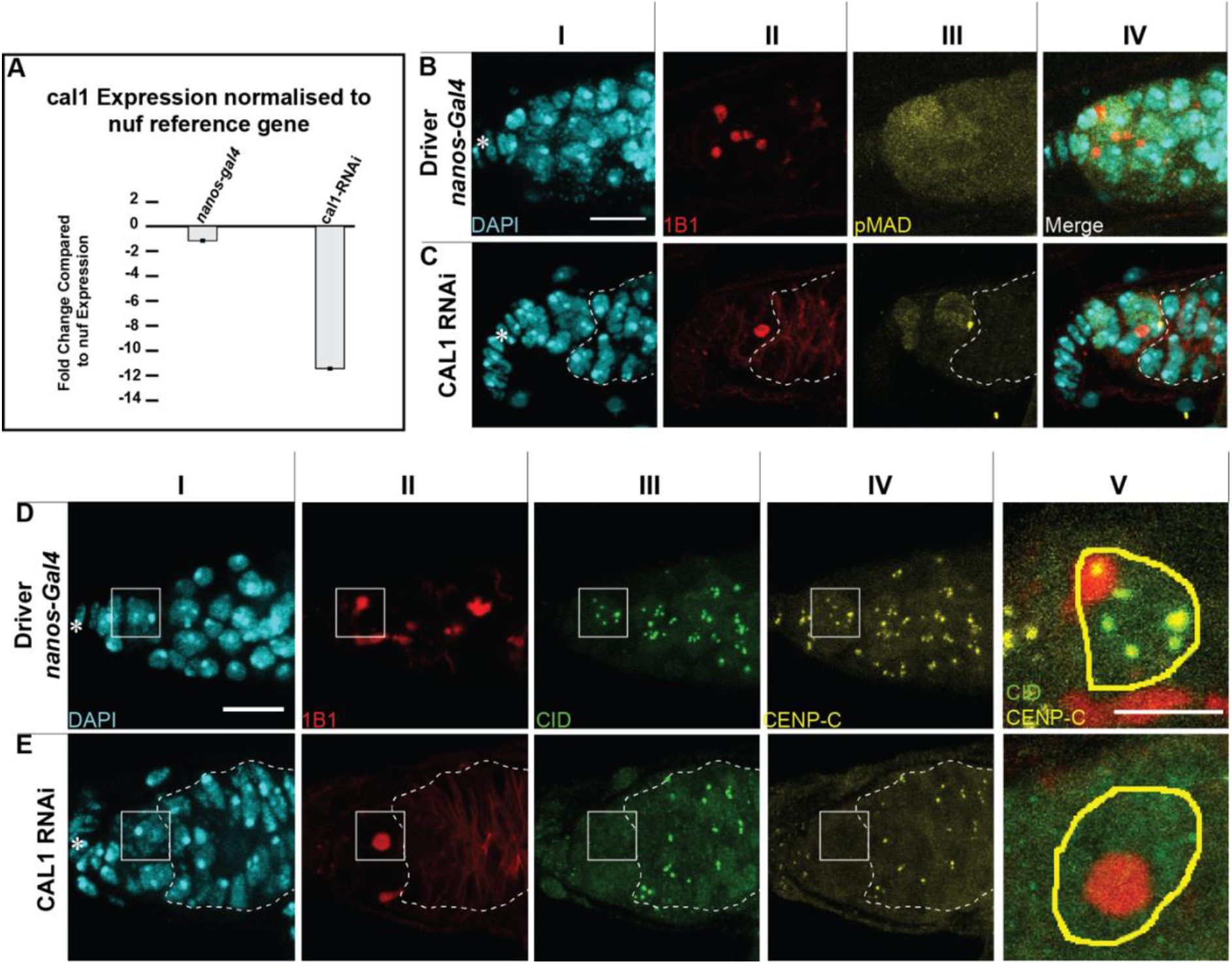
CAL1 knock down blocks cell proliferation. **A**) CAL1 knock down confirmation by real time-qPCR. **B-C**) Confocal z-stack projection of a germarium of *nanos-Gal4* (**B**) and CAL1 RNAi (**C**) flies, stained for DAPI (cyan), anti-1B1 (spectrosome, red) and anti-pMAD (labels GSCs, yellow). **D-E**) Confocal z-stack projection of a germarium of *nanos-Gal4* (**D**) and CAL1 RNAi flies (**E**), stained for DAPI (cyan), anti-1B1 (spectrosome, red), anti-CID (green) and anti-CENP-C (yellow). Star indicates the position of the terminal filament; dotted lines represent follicle cells; 3 days old female flies; scale bar: 10 μm; for **DV and EV** scale bar: 5 μm.

### Centromeric CAL1 is required for CID and CENP-C recruitment to GSC centromeres

Given that CAL1 is located both at the centromeres and in the nucleolus, we investigated the depletion of both CAL1 pools in GSCs. We performed antibody staining against centromeric and nucleolar components in combination with CAL1 (Fig. S3C). In *nanos-Gal4* flies (Fig. S3C), CAL1 co-localises with both CENP-C and the nucleolar marker FIBRILLARIN respectively (Fig. S3C and S3C). In the CAL1-knocked down samples, CAL1 is still present in the nucleolus of GSCs, as it co-localises with FIBRILLARIN (Fig. S3C), however it is missing from the centromeres, as we could not distinguish any CAL1 signal outside of the nucleolus overlapping with the CENP-C marker (Fig. S3C). Indeed, neither CENP-C nor CID is detectable at centromeres in the knocked down cells (Fig. 4D-E). From our observation we conclude that: i) our knock down depletes the pool of CAL1 proteins at the centromeres, but not the one at the nucleolus; ii) knock down of centromeric CAL1 is responsible for the loss of functional centromeres in GSCs.

### Over-expression of CAL1 and CID promotes stem cell self-renewal, while CAL1 over-expression promotes cell proliferation

To further explore centromere function in GSCs, we performed an over-expression of CAL1 together with CID, and CAL1 over-expression alone. For this purpose, we generated transgenic flies carrying CAL1-YFP only, or both CAL1-YFP and CID-mCherry, all under the control of UASp (see STAR methods). We then crossed both lines to a *nanos-Gal4* driver line. Ovaries from F1 progeny were dissected and first screened for correct localisation of the fusion proteins, confirmed using antibody staining against CID and FIBRILLARIN (Fig. S4). As expected, the *nanos-Gal4* control does not show any YFP or RFP fluorescence (Fig. S4A-E’). In the CAL1-CID over-expression, CAL1-YFP co-localises with CID-mCherry and with CID antibody (Fig.S4F-I stylised arrow and inset S4J), but we could not detect any co-localisation with FIBRILLARIN in the nucleolus (Fig. S4F’-I’ stylised arrow and inset S4J’). In the CAL1-YFP over-expression, instead, CAL1 localises as expected, at centromeres (Fig. S4K-N stylised arrows and inset S4O) and at the nucleolus (Fig. S4K’-N’ arrow heads and inset S4O’). We next verified any change in the niche composition using antibodies against GSC and CB markers. We first quantified the number of spectrosomes (Fig. 5A-C). In *nanos-Gal4* we found an average of 2 spectrosomes/germarium (Fig. 5A arrows, and 5J; nanos-Gal4=1.84±0.16, n=30 germaria). In the CAL1-CID over-expression, this number increases about 1.4 fold (Fig. 5B arrows, 5J; UAS_CAL1-UAS-CID=2.61±0.17, n=30 germaria), while in the CAL1 over-expression, the number of spectrosomes/germaria almost doubles (Fig. 5C arrows, 5J; UAS_CAL1=3.51±0.31, n=30 germaria). To better understand the disruption caused to the niche, we used the additional markers pMAD, a stem cell marker (Song, 2004) and SEX-LETHAL (SL) (Fig. 4I-N), a marker that labels the GSC-CB transition. SL is present from GSCs up to the 2-cell cyst stage and can therefore be used to define the size of the niche (Chau et al., 2009). In this case our quantification shows that *nanos-Gal4* germaria have approximately one pMAD positive cell (*nanos-Gal4*_pMAD_=1.13±0.14, n=30 germaria; Fig. 5D and 5K), while the number of pMAD positive cells doubles in both over-expression lines (Fig. 5E-F and 5K; UAS_CAL1-UAS_CID_pMAD_=2.10±0.13; UAS_CAL1_pMAD_=2.20±0.13). Antibody staining against SL revealed that there is no difference in the size of the niche between the *nanos-Gal4* control and the CAL1-CID over-expression (*nanos-Gal4*_SL_=4.83±0.32, n=30 germaria; UAS_CAL1-UAS_CID_SL_=4.86±0.25; Fig. 5G-H and 5K), while over-expression of CAL1 alone is responsible for an increase in the size of the niche compared to the control (UAS_CAL1_SL_=6.86±0.27, n=30 germaria; Fig. 5I and 5K). This analysis shows that both CAL1-CID over-expression and CAL1 over-expression increase the number of GSCs compared to the control *nanos-Gal4*. Furthermore, over-expression of CAL1 alone is responsible for an increase of the number of germ cells present in the germarium.

**Figure 5.**
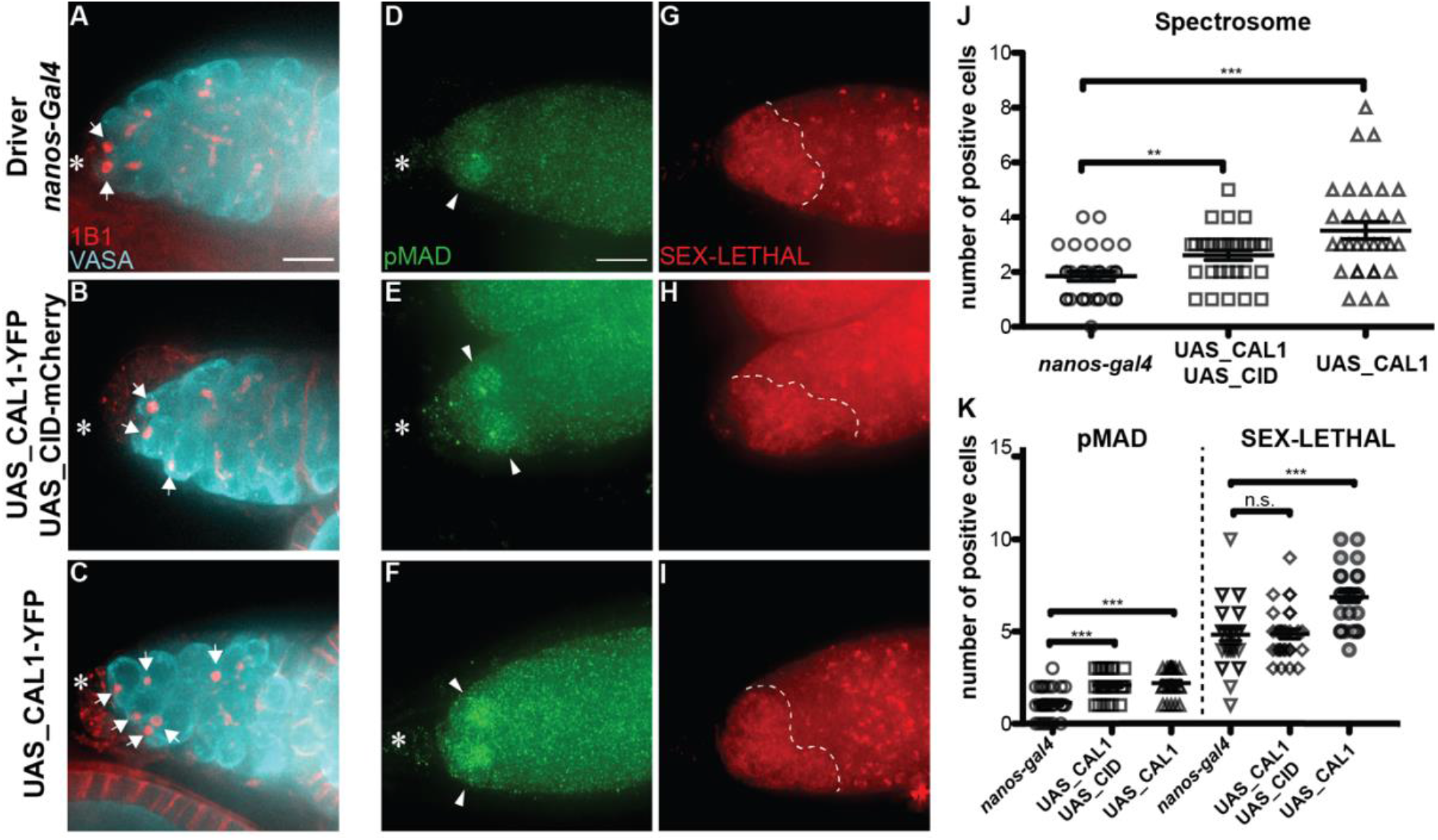
CID and CAL1 over-expression promote stem cell self-renewal. **A-C**) widefield z-stack projection of a germarium of *nanos-Gal4* (**A**), UAS_CAL1-YFP_UAS_CID-mCherry (**B**) and UAS_CAL1-YFP (**C**) flies, stained for DAPI (cyan) and anti-1B1 (spectrosome, red). **D-I**) Confocal z-stack projection of a germarium of *nanos-Gal4* (**D** and **G**), UAS_CAL1-YFP_UAS_CID-mCherry (**E** and **H**) and UAS_CAL1-YFP (**F** and **I**) flies, stained for anti-pMAD (green) and anti-SEX-LETHAL (red). **J**) Spectrosome quantification. **K**) pMAD and SEX-LETHAL quantification. Data are represented as the mean±standard error of the mean (SEM); ***= p value<0.0005; **= p value< 0.005. Star indicates the position of the terminal filament; arrows indicate the spectrosome; arrow heads indicate pMAD positive cells; dotted lines represent SEX-LETHAL region; 3 days old female flies; scale bar: 10 μm.

When expressed as a SL/pMAD ratio (Fig. S4P), the ratio does not change between the control *nanos-Gal4* and CAL1-YFP over-expression (*nanos-Gal4*_SL/pMAD_=3.57±0.32; UAS_CAL1_SL/pMAD_=3.38±0.22, Fig. S4P). This is the result of an increase of both GSCs, as well as an increase in the size of the niche in the germaria over-expressing CAL1. However, in the CAL1-CID over-expression this ratio is significantly lower compared to the control (UAS_CAL1-UAS_CID_SL/pMAD_= 2.47±0.23, Fig. S4P). This is because an increase in GSCs number it does not correspond an increase in the size of the niche. This means that GSCs in germaria over-expressing CAL1-CID self-renew, rather than differentiate. To exclude the possibility that phenotypes were due to the genetic background of the responder flies, we conducted the same analysis on the ovaries of female flies of the transgenic responder lines (in which the over-expression is not induced) and similar values to the *nanos-Gal4* control (not shown). Altogether, our results suggest that the over-expression of both CAL1 and CID together promotes self-renewal, while the over-expression of CAL1 alone stimulates proliferation.

### CAL1 function is dispensable in the 16-cell cysts egg chamber

To examine CAL1 and CID requirements at later stages of egg development, we performed knock down experiments in cysts outside of the stem cell niche using the *bam-Gal4* driver (active in 4-8 cell cysts). Female flies from the control and the knock down were dissected and stained for VASA to mark germ cells and BAM to mark 4-8 cell cysts. Surprisingly, we noticed that cell division past the 4-8 cell stage is not impaired by depletion of either CID or CAL1 (Fig. 6A-I’). Specifically, in the CID RNAi we did not observe a significant diminishment of CID levels at the centromeres, compared to the control (Fig. 6J-K). In the CAL1 RNAi, CID levels appear to be reduced, but cell division proceeds normally (Fig. 6L). In 16-cell cysts, after the BAM region, we confirmed a reduction of CAL1 in the CAL1-knocked down samples, compared to the *bam-Gal4* drivers (Fig. S5A-H’). In order to check the impact of this reduction on centromere assembly, we antibody stained samples against CENP-C. In the control cysts, identified through the fusome morphology (Figure 6M-P and inset 6M’-P’, arrow in inset Fig. 6N’), 2-4 centromere foci closely opposed to the nucleolus are normally visible (Unhavaithaya and Orr-Weaver, 2013) (Fig. 6O-P and inset Fig. 6O’-P’). In the CAL1 knocked down sample (Fig. 6Q-T and inset Fig. 5Q’T’), we did not observe any change in the amount of CENP-C (Fig. 6S-T and inset 6S’-T’). These results indicate that CID and CENP-C are already assembled at centromeres at this stage and CAL1 function is dispensable at least for the cell division occurring after the 8-cell stage.

**Figure 6.**
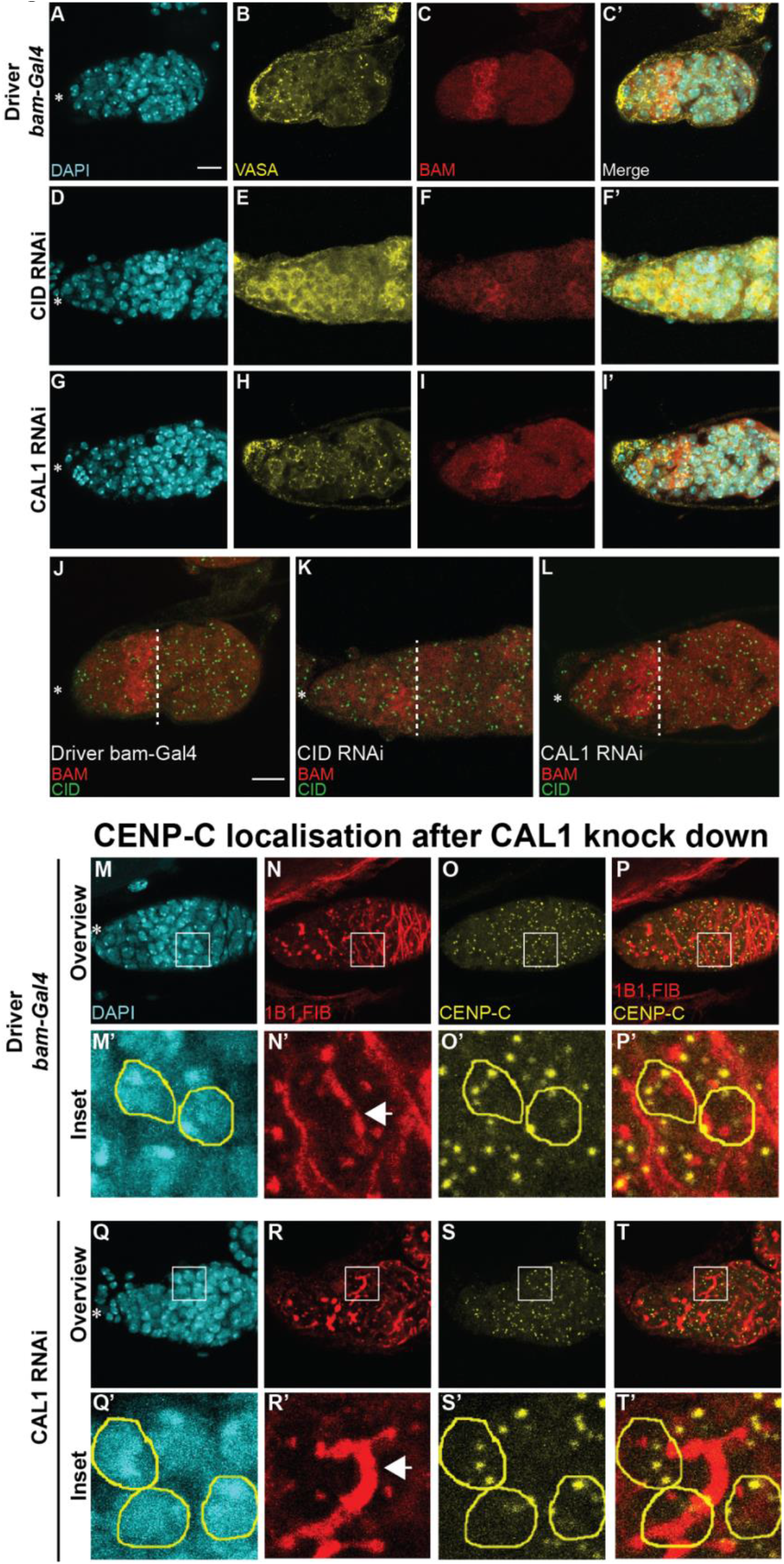
CAL1 role at centromeres is dispensable in the 16 cell cysts egg chamber. **A-I’**) Confocal z-stack projection of a *Drosophila* germarium of *bam-Gal4* (**A-C’**), CID RNAi (**D-F’**) and CAL1 RNAi (**G-I’**) flies, stained for DAPI (blue), anti-VASA (yellow) and anti-BAM (red). **J-L**) Confocal z-stack projection of a *Drosophila* germarium of *bam-Gal4* (**J**), CID RNAi (**K**) and CAL1 RNAi (**L**) flies, stained for anti-BAM (red) and anti-CID (green). **M-T’**) Confocal z-stack projection of a germarium of *bam-Gal4* (**M-P** and inset **M’-P’**) and CAL1 RNAi (**O-T** and inset **O’-T’**) flies, stained for DAPI (blue), anti-1B1 (red), anti-FIBRILLARIN (red) and anti-CENP-C (yellow). 16-cell cysts were selected based on the fusome morphology (arrow) in the control (**M’-P’**) and in the CAL1 RNAi (**O’-T’**). Star indicates the position of the terminal filament; dotted line indicates the end of the BAM positive region; arrow indicates the fusome; 3 days old female flies; scale bar: 10 μm; inset scale bar: 5 μm.

### CID assembly dynamics differ between GSCs and differentiated cells outside of the niche

To better understand the dynamics of CID assembly between GSCs and cysts, we measured the amount of CID per nucleus in both cell types to detect possible differences. Ovaries from *nos-Gal4* flies were dissected and antibody-stained against CID and we used the marker H3S10P to specifically capture cells at prophase. Indeed, prophase is the cell cycle timing at which centromere assembly is complete in GSCs and it can also be used to mark cysts at the 8-cell stage, as CCs undergo synchronous mitoses (Fig. 7A-H). We noted that anti-CID staining at prophase labels centromeric CID, but also shows a nuclear non-centromeric localisation. As we did not observe this localisation with CID-GFP, it is possible that it results from this specific antibody combination. To overcome this, we focused our quantification on centromeric CID signals only. Compared to prophase GSCs, cyst nuclei are smaller yet centromeric foci are present in a similar number to GSCs at this stage. From our quantification, we detected an approximate 40% diminishment of the total amount of CID at the 8-cell stage (CC=323.4±20.94, n=26 cell analysed; Fig7I), compared to GSCs (GSC=547.2±41.57, n=24 cell analysed; Fig.7I). This indicates a dramatic change in the assembly dynamics of CID into the centromeric chromatin, such that it is not replenished to 100% each cell division. Because we did not observe any significant reduction in CID amount either using CID RNAi or CAL1 RNAi (Fig. 6 and S5), these data suggest the inheritance of CID from the niche together with the reduced new CID loading in these cells.

**Figure 7.**
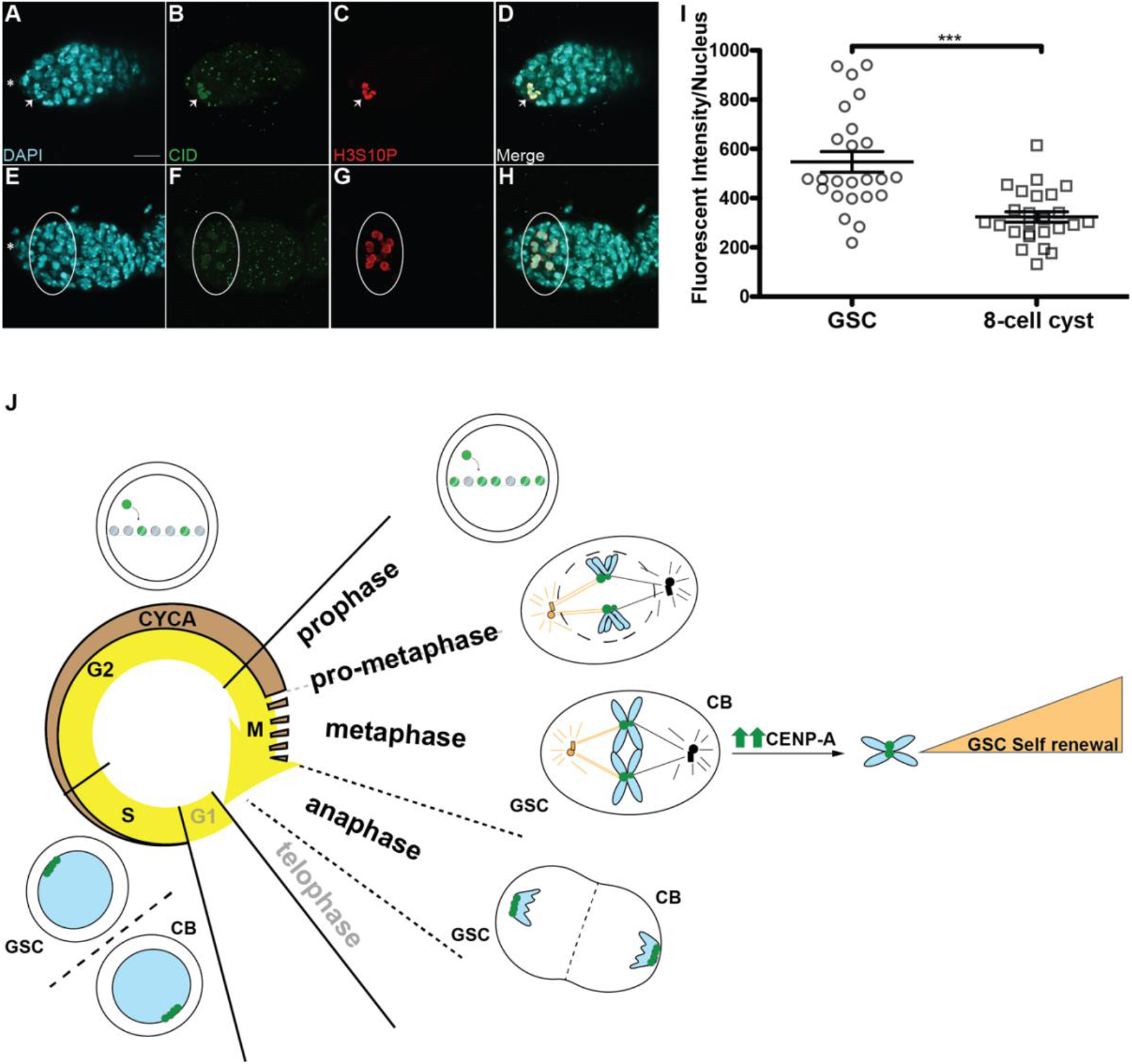
Cysts outside of the niche incorporate less CID compared to GSCs. **A-H**) Confocal z-stack projection of a germarium of *nanos-Gal4*, stained for DAPI (cyan), anti-CID (green) and anti-H3S10P, to highlight GSC (**A-D**, arrow) and 8 cell cysts (**E-H**, circle) in prophase. **I**) Quantification of fluorescence intensity of CID at centromeres in GSCs and 8 cell cysts at prophase obtained using widefield microscopy. Fluorescence Intensity is expressed as integrated density after background subtraction (see material and methods). Data are represented as the mean±standard error of the mean (SEM); ***= p value<0.0001. Star indicates the position of the terminal filament; 3 days old female flies; scale bar: 10 μm **J**) Model for centromere assembly during the cell cycle. During G2 phase, centromere assembly starts, promoted by CYCA, and centromeric nucleosomes (green) replace canonical nucleosomes (grey) at the centromeres. This process continues until prophase. At prometaphase the nuclear envelope breaks down (dotted lines) and microtubules from the centrosomes attach centromeres through the kinetochore. At this point sister chromatid pairs have been loaded with differential amount of CID at centromeres. Chromosomes that retain the higher amount of CID (bigger centromeres, figurative) are able to attract more nucleating from the daughter centrosome (orange) and that will be inherited by the GSC. At anaphase centromeres are clustered at the opposite sides of the two daughter nuclei and strikingly retain this localisation also during replication. At these stages we did not detect any changes in the amount of CID between GSC and CB. Telophase and G1 phase are shown as transparent because of the lack of data for these two phases. Increasing CID expression can determine cell fate, promoting GSC self-renewal.

## DISCUSSION

Functional centromeres are fundamental for a correct segregation of chromosomes during cell division. In this study we have performed a detailed characterisation of centromere dynamics throughout the cell cycle in *Drosophila* female GSCs. This analysis reveals that GSCs initiate CID incorporation during G2 phase and that its deposition continues until prophase (Fig. 7J) and *Drosophila* neural stem cells follow the same trend. Notably, this timing is different from existing studies in other metazoans. We also found that CYCA, a G2 phase regulator, is critically involved in CID (and CENP-C) loading at centromeres, acting as a new centromeric assembly factor possibly independent of CAL1 (Fig. 7J). Moreover, chromosomes that will be inherited by GSCs are labelled with higher amount of CID (Fig. 7J), detectable at prometaphase and metaphase. We show for the first time that depletion of the centromeric protein CAL1 in the stem cells is responsible for the lack of differentiated cells, but not of stem cells, in *Drosophila* germline female tissues. Moreover, over-expression of CAL1 and CID together promotes stem cell self-renewal, while CAL1 over-expression alone causes GSC-like tumours. Intriguingly, we find that CAL1 function becomes dispensable at later stages of germline differentiation, probably as CID is incorporated in GSCs and inherited by cells outside of the niche. According to our findings we raise three main points of discussion:

### 1) Biological significance of centromere assembly in G2-M phase

a. ***Cell cycle time***. The assembly of centromeres between G2 and prophase, supported by our quantification and functional analysis (Fig. 1 and 2), could be due to the contraction of the G1 phase, as a characteristic of stem cells (Becker et al., 2006; Pauklin and Vallier, 2013; White and Dalton, 2005). Given that G1 phase is missing in the fly embryonic divisions and CID loading occurs at anaphase (Schuh et al., 2007), G2-M assembly might be a unique property of stem cells. Remarkably, this timing is similar to the one found for *Drosophila* spermatocytes, which assemble centromeres in prophase of meiosis I (Dunleavy et al., 2012; Raychaudhuri et al., 2012). These spermatocytes undergo an arrest in prophase I for days, indicating a gradual loading of CID over a long period of time. Intriguingly, a similar phenomenon has been recently observed in G0-arrested human tissue culture cells and prophase I-arrested starfish oocytes (Swartz et al., 2018). A possibility is that cells delayed in a specific cell cycle phase have developed mechanisms to actively maintain centromeres by a slow but constant deposition of CENP-A (Swartz et al., 2018). In this context, *Drosophila* GSCs biology and our results well support this hypothesis. Indeed, GSCs divide approximately once every 24 hours (Yamashita, 2003) and spend the majority of their time in G2 phase. Furthermore, according to our data centromere assembly occupies a wide window of time from G2 phase to prophase. Therefore, *Drosophila* GSCs might show similar properties of quiescent cells.
b. ***Cell cycle regulation***. Incorporation of CID prior to chromosome segregation might reflect a different CYCLIN-CDK activity in these cells. For instance, it has been already shown that in *Drosophila* GSCs CYCLIN E, a canonical G1/S cyclin, exists in its active form (in combination with Cdk2) throughout the cell cycle, indicating that some of the biological process commonly occurring in G1 phase, might actually take place in G2 phase (Ables and Drummond-Barbosa, 2013). This is in line with our functional findings, where depletion of CYCA causes a decreased efficiency in the assembly of centromeres, measured as a loss of CID and CENP-C at centromeres.
c. ***Epigenetic mechanism to drive cell fate during stem cell asymmetric division***. Centromere assembly in G2 phase can also represent a strategy to differentially label sets of chromosomes to avoid random segregation during asymmetric division of the stem cell. In this context, we show that sister chromatids are labelled with differential amounts of CID upon centromere assembly; chromosomes that will be inherited by the GSCs incorporate 20% more CID, compared to chromosomes that will be inherited by the CBs (Fig. 3). Specifically, we observed fluctuation of CID levels between sister centromere pairs during GSC prometaphase/metaphase, as already described (Bodor et al., 2014). Despite this, a clear asymmetry is observed between the total amount of CID present on the set of chromosomes belonging to “GSC side” versus the one of the “CB side”. More importantly, this asymmetry is not observed in CCs during the same time window. Moreover, arrested starfish oocytes and quiescent human tissue-cultured cells are able to incorporate small amount of CENP-A per day (in the order of 2-10%), which is has an effect on centromere assembly in the long run (Swartz et al., 2018). Furthermore, this small excess of CID is used to capture more microtubules nucleating from the daughter centrosome during the G2-M transition, the time at which centromere assembly occurs according to our data (Fig. 7J). Finally, our results are in line with the general concept of centromere assembly, which always occurs in a defined period of the cell cycle, outside of replication where canonical histones are incorporated into the chromatin.

### 2) Novel role for CID in cell fate during stem cell asymmetric division

Our functional knockdown and over-expression studies support a role for CAL1 in cell proliferation (Fig. 4 and 5). Indeed, centromeric proteins have been already proposed as bio-markers for cell proliferation (Swartz et al., 2018). In parallel, we show that the over-expression of both CID and CAL1 increases the number of GSCs without increasing the size of the niche, promoting stem cell self-renewal (Fig. 5). As we did not observe CAL1 localisation to the nucleolus in germaria over-expressing both CAL1 and CID, our data suggest that this phenotype is strongly linked to the centromere locus. Finally, our quantification and functional analysis together support the hypothesis according to which amount of CENP-A detected at the centromeres might drive stem cell identity during stem cell mitosis.

### 3) Centromere assembly occurs prior to meiosis

Our quantitative analysis of total CID levels in GSCs and downstream cyst cells, as well as CAL1 and CID knock down after the 8-cell stage of the germarium (Fig. 6 and 7) indicate that centromeres are crucially assembled in the GSCs and therefore before meiosis of the oocyte takes place. Thus, it is possible that the 16-cell cysts inherit centromeric proteins synthesized and deposited in the GSCs and little new CID loading occurs. This would explain why CAL1 function at centromeres is dispensable in these cells.

Ultimately, our results provide the first functional evidence that centromeres have a role in the epigenetic pathway that specify stem cell identity. Furthermore, our data support the silent sister hypothesis according to which centromeres can drive stem asymmetric division.

## Acknowledgments

E.M.D., A.M.K. and A.E.W. are funded by Science Foundation Ireland -PIYRA 13/YI/2187. A.A.D. is funded by Government of Ireland Postdoctoral Fellowship 2017/1324 and Science Foundation Ireland – PIYRA 13/YI/2187. B.L.C. is funded by Government of Ireland Postgraduate Fellowship 2018/1208. The authors acknowledge the facilities and technical assistance of the NCBES qPCR Facility, funded by NUIG and the Irish Government’s Programme for Research in Third Level Institutions, Cycles 4 and 5, National Development Plan 2007-2013, the Centre for Microscopy & Imaging at the National University of Ireland Galway (www.imaging.nuigalway.ie) and the Edinburgh Super-Resolution Imaging Consortium at the Institute of Genetics and Molecular Medicine at the University of Edinburgh (https://www.esric.org/). Stocks were obtained from the Bloomington Drosophila Stock Center (NIH P40OD018537). Antibodies obtained from the Developmental Studies Hybridoma Bank, created by the NICHD of the NIH are maintained at The University of Iowa, Department of Biology, Iowa City, IA 52242. We thank Gary Karpen and Sylvia Erhardt for antibodies.

## Author contribution

A.A.D. and E.M.D. conceived and designed the study. A.A.D. performed the molecular biological experiments, fluorescence microscopy imaging and data analysis in the experiments presented in Fig. 1–2–3–4–6. B.L.C. performed the molecular biological experiments, fluorescence microscopy imaging and data analysis in the experiment presented in Fig. 7A-I. A.A.D and A.M.K. performed the molecular biological experiments, fluorescence microscopy imaging and data analysis in the experiments presented in Fig. 5. A.A.D and A.W. generated the transgenic fly lines. A.A.D. and E.M.D. wrote the manuscript.

## STAR METHODS

### Generation of the transgenic flies

Transgenic lines expressing C terminal tagged CAL1-YFP or both CAL1-YFP-CID-mCherry under control of UASp sequences were generated by transposable (P) element transformation of pUASp vector (kind gift from Xin Chen) in w^1118^embryos (injection, selection and balancing by BestGene Inc). Specifically, CAL1-YFP and CID-mCherry constructs were placed in tandem following UASp sequences in the same plasmid. *cid* and *cal1 cDNA* were amplified from wild type. mCherry containing three codons for glycine residues at both sides was inserted in between *cid N-terminal* and *cid C-terminal* as described (Schuh et al., 2007).

### Fly stocks and husbandry

Stocks were cultured on standard cornmeal medium (NUTRI-fly) preserved with 0.5% propionic acid and 0.1% Tegosept at 20°C under a 12 hours light-dark cycle. *uas*-RNAi lines were obtained from the Bloomington Stock Centre (CYCA #35694) and Vienna Drosophila RNAi Centre (CAL1 #45248; CENP-A/CID #102090). The germline-specific promoters *nanos* and *bam* were used to drive GAL4 expression (P{w[+mC]=UAS-Dcr-2.D}1, w[1118]; P{w[+mC]=GAL4-nos.NGT}40, provided by Bloomington Stock Centre, #25751; *bam-Gal4* was a kind gift from Margaret T. Fuller).

Crosses were performed at 20°C, 25°C and 29°C, specifically: CAL1 and CID knock down using the *nanos-Gal4* driver were carried out both at 25 and 20 °C, while CAL1 and CID knock down using the *bam-Gal4* driver were conducted both at 29 °C. CYCA knock down with *nanos-Gal4* driver was performed at 25 °C. Crosses for over-expression were carried out using *nanos-Gal4* driver at 25°C. Transgenic lines expressing GFP-tagged CENP-A/CID and RFP-tagged H2Av (heterozygotes) (Schuh et al., 2007) under respective endogenous promoters were a kind gift from Christian Lehner. Results obtained from each experiment rely on three biological replicates, unless otherwise specified.

### Immunofluorescence (IF)

GSCs usually undergo mitotic division at very low frequency (Yamashita, 2003), therefore, to increase our chance to catch multiple cell cycle phases during cell division at once, we used young female flies (<1 day old) for centromere assembly quantifications. To quantify metaphase GSCs in Fig. 3 we used young female flies 30 min old. For all the other experiments we used 3 days old female flies. Ovaries from young adult females were dissected in 1XPBS and fixed in 4% paraformaldehyde. In order to carry out the tubulin staining, ovaries were fixed ice cold methanol for 20 min at 4°C, followed by Acethone at −20 °C for additional 2 min. After fixation, samples were immediately washed in 1XPBS-0.4%Triton-X100 (0.4PBT). Samples were then blocked in 0.4PBT with 1% BSA for 3-4 hours at room temperature, incubated with primary antibodies (in blocking buffer) overnight at 4°C and with secondary antibodies (in blocking buffer) for 1 hour at room temperature. For quantification of CID in neuroblasts, brains from 3^rd^ instar larvae were dissected and fixed as described above.

### EdU Assays

Young female flies were dissected and incubated for 30 min with EdU (0.01 mM) in PBS 1x and then fixed as described. After washing in 0.4PBT, ovaries were incubated for 30 min in the dark with 2 mM CuSO4, 1–100 M fluorescent azide (from 10 to 100mM stocks in DMSO), and 10 mM ascorbic acid. Samples were then washed with 0.4PBT for 10 min and then blocked and stained as previously described.

### Antibodies

For immunostaining, the following antibodies were used: rabbit anti-CENP-A (CID) antibody (Active Motif 39719; 1:500), rat anti-CID (Active Motif 61735; 1:500), guinea pig anti-CENP-C (Erhardt et al., 2008) (1:1000), rabbit anti-H3S10P (Abcam ab5176, 1:1000), mouse anti-H3S10P (Abcam ab14955; 1:1000), rabbit anti-H3T3P (MERK 05-746R; 1:1000), rabbit anti-VASA (Santa Cruz sc-30210; 1:250), goat anti-VASA (Santa Cruz sc-26877; 1:100), mouse anti-Fibrillarin (Abcam ab4566; 1:500), mouse anti-BAM (DSHB ab10570327; 1:10), mouse anti-CYCA (DSHB, A12 ab528188; 1:250), rabbit anti-pMAD (Abcam ab52903; 1:250), mouse anti-SPECTROSOME/1B1 (DSHB ab528070; 1:50), rat anti-DEADPAN (Abcam 195173; 1:100), mouse anti-tubulin (Abcam ab44928; 1:100) and rabbit anti-CAL1 (Bade et al., 2014) (1:1000).

### Confocal microscopy

Images of immunostained ovaries were taken using an inverted Fluoview 1000 laser scanning microscope (Olympus) equipped with a 60x oil immersion UPlanS-Apo objective (NA 1.2). The samples were excited at 404, 473, 559, 635 nm respectively for DAPI, Alexa Fluor 488, 546 and 647. Light was guided via D405/473/559/635 dichroic mirror (Chroma) to the sample. The emission light was guided via a size adjustable pinhole, set at 115 μm. Fluorescence passed through a 430-455nm; 490-540-nm; 575-620-nm; 655-755 nm band-pass filter for detection of respectively DAPI, Alexa Fluor 488, 546 and 647 in sequential mode. Images were acquired as z-stacks with a step size of 0.5 μm.

### Super-resolution microscopy

Super-resolution images of immunostained ovaries were acquired using structured illumination microscopy. Samples were prepared on high precision cover-glass (Zeiss, Germany). 3D SIM images were acquired on a N-SIM (Nikon Instruments, UK) using a 100x 1.49NA lens and refractive index matched immersion oil (Nikon Instruments). Samples were imaged using a Nikon Plan Apo TIRF objective (NA 1.49, oil immersion) and an Andor DU-897X-5254 camera using 405, 488, 561 and 640nm laser lines. Z-step size for Z stacks was set to 0.120 um as required by manufacturers software. For each focal plane, 15 images (5 phases, 3 angles) were captured with the NIS-Elements software. SIM image processing, reconstruction and analysis were carried out using the N-SIM module of the NIS-Element Advanced Research software. Images were checked for artefacts using the SIM check software (http://www.micron.ox.ac.uk/software/SIMCheck.php). Images were reconstructed using NiS Elements software v4.6 (Nikon Instruments, Japan) from a z stack comprising of no less than 1um of optical sections. In all SIM image reconstructions the Wiener and Apodization filter parameters were kept constant.

### Widefield microscopy

Images were acquired using a DeltaVision Elite microscope system (Applied Precision) equipped with a 100x oil immersion UPlanS-Apo objective (NA 1.4). Images were acquired as z-stacks with a step size of 0.2 μm. Fluorescence passed through a 435/48 nm; 525/48 nm; 597/45 nm; 632/34 nm band-pass filter for detection of respectively DAPI, Alexa Fluor 488, mCherry and Alexa fluor 647 in sequential mode.

### Quantification

For each quantification one cell/germarium was considered, unless differently specified. Images from a single cell were projected (max intensity) to capture all the centromeres present in the cell at a specific cell cycle phase. Image J software (Schneider et al., 2012) was used to measure fluorescent intensity of CID in the following way. The background was subtracted from the projected image. Threshold was adjusted and the image was converted to binary. Overlapping centromeres were separated using the command “watershed”. Following, the command “analyse particles” was used to select centromeres. Size was adjusted, in order to eliminate unwanted objects. Finally, mean grey values (MGV) from each centromere foci were extracted and used as fluorescent intensity to measure the total amount of fluorescence per nucleus and or per individual foci. For stages such as replication, metaphase and anaphase, in which centromeres are clustered, it was necessary to analyse each individual centromere Z-stack-slice by slice. In the quantification of CID in neuroblasts and in metaphasic GSCs we used the integrated density (MGV*Area) instead of the mean grey values, due to the strong clustering observed after projection. In the quantification of CID in GSC and cysts we used the integrated density (MGV*Area) instead of the mean grey values, due to the strong clustering observed after projection in the cysts. Quantification of pMAD and SEX-LETHAL positive cells was obtained counting the positive cells for each signal through the z-stack of each image. Statistical analysis was performed using prism software.

### RNA Isolation and Quantitative PCR

Wild type, drivers (*nanos-Gal4*) and CAL1 knocked down flies were collected 7 days after hatching at 20°C and then dissected to extract ovaries. Total RNA from each sample was stored in TRIzol (1559-6026, Ambion, Life Technologies) at −80°C until processed. RNA extraction and purification was carried out with Rneasy MinElute clean up kit (Cat# 74204, Qiagen). All the isolated RNAs were then standardised to the same concentration and cDNA was synthesized (4387406, High Capacity RNA to cDNA, Thermofisher-Biosciences). Quantitative PCR was performed using the Applied Biosystem StepOnePlus Instrument and Power Up Sybr Green Master mix (Cat# A25780, Applied Biosystem). The following genes, from *Drosophila* genome, were considered for this experiment: *Glyceraldehyde 3-phosphate dehydrogenase (gapdh); Ribosomal Protein L32 (rpl32/rp49); nuclear-fall-out (nuf); cal1*. Primers for all the considered genes were designed using MacVector (www.macvector.com) to amplify 75-150 base-pair fragments of the desired gene (Fig. S2A). Prior the qPCR experiment, these primers were tested with a mixture of cDNA from *Drosophila* ovaries to make sure they only will amplify a single region from the genome. Following, we checked primer efficiencies with a dilution curve (10^−1^-10^−5^) to make sure their range was within the negligible value of 1.9-2.0. Among the reference genes considered (*gapdh; rpl32/rp49; nuf*), *nuf* (*nuclear-fall-out*) is highly stable at 20 °C, between both control and the knocked down sample. Therefore, qPCR samples were standardised with *nuf* and relative fold change values were calculated in Microsoft Excel and were standardized against our reference gene based on published formulas (Livak, 2001). Each qPCR experiment consists of two biological replicates and each sample was analysed using three technical replicates per qPCR experiment. According to the MIQE guidelines (Bustin et al., 2009), primer sequences used, relative efficiency and amplification factor used in the calculation together with the melt curve obtained are provided: *gapdh*, Fw-GCTGGTGCCGAATACATCGTGG; Rv-CCAAGTTGACGCCGCAAACG; efficiency: 90.7%, amplification factor:1.91; *rpl32/rpl49*, Fw-CCGCTTCAAGGGACAGTATCTGATGC; Rv-TTCTGCATGAGCAGGACCTCCAGC; efficiency:88.1%; amplification factor 1.89; *nuf*, Fw-TGGCGAAAATGAGTATCCCACCC; Rv-GGTTGTGTCCACTGTTGTTACCCACG; efficiency: 105.8%; amplification factor: 2.06; *cal1*, Fw-GTGAACGACAAGAGATTCCAGCGAC; Rv-AGTCCCTGCTCGGTCAGTGTGAAG; efficiency:102.9%; amplification factor: 2.03.

## SUPPLEMENTAL INFORMATION

**Figure S1. CYCLIN A knock down confirmation, related to Figure 2. A-F**) Confocal z-stack projection of a *Drosophila* germarium of *nanos-Gal4* (**A-C**) and CYCA RNAi (**D-F**) flies at 25 °C, stained for DAPI (blue), anti-CID (red) and anti-CYCA (green); scale bar: 10 μm.

**Figure S2. GSCs in prometaphase and metaphase used for quantification of CID present on the GSC side and CB side, related to Figure 3. A-N**) Super resolution (N-SIM) z-stack projection of a *Drosophila* GSC of a *wild type* germarium, stained for DAPI (blue), anti-CID (red), anti-H3T3P (green) and anti-SPECTROSOME (magenta); scale bar: 5 μm; inset scale bar: 1 μm.

**Figure S3. CAL1 is required for a correct development of the egg in *Drosophila* germline tissues, related to Figure 4. A**) Confocal z-stack projection of a *Drosophila* germarium of *nanos-Gal4*, CID RNAi and CAL1 RNAi flies, stained for DAPI (blue), anti-VASA (yellow) and anti-1B1 (spectrosome, red). **B**) Confocal z-stack projection of a *Drosophila* germarium of *bam-Gal4*, CID RNAi and CAL1 RNAi flies, stained for DAPI (blue), anti-VASA (yellow) and anti-BAM (red). **C**) Confocal z-stack projection of a *Drosophila* germarium of *nanos-Gal4* (20 °C) stained for DAPI (blue) and anti-VASA (yellow), anti-FIBRILLARIN (magenta) and anti-CAL1 (green) and stained for DAPI (blue) and anti-1B1 (red), anti-CENP-C (yellow) and anti-CAL1 (green). Confocal z-stack projection of a CAL1 RNAi (20 °C) stained for DAPI (blue) and anti-VASA (yellow), anti-FIBRILLARIN (magenta) and anti-CAL1 (green) and stained for DAPI (blue) and anti-1B1 (red), anti-CENP-C (yellow) and anti-CAL1 (green). Star indicates the position of the terminal filament; 3 days old female flies; scale bar: 10 μm.

**Figure S4. Over-expression of both CAL1 and CID promotes GSC self-renewal, related to Figure 5. A-O’**) Confocal z-stack projection of a *Drosophila* germarium of *nanos-Gal4*, UAS_CAL1-YFP_UAS_CID-mCherry and UAS-CAL1-YFP flies, stained for anti-CID and or anti-FIB (cyan). Star indicates the position of the terminal filament; arrows indicates the centromeres; arrow heads indicate the nucleolus; 3 days old female flies; **P**) Ratio of the number of SEX-LETHAL positive cells/number of pMAD positive cells. scale bar: 10 μm; inset scale bar: 5 μm.

**Figure S5. CAL1 knockdown confirmation using the *bam-Gal4* driver, related to Figure 6**. Confocal z-stack projection of a germarium of *bam-Gal4* (**A-D** and inset **A’-D’**) and CAL1 RNAi (**E-H** and inset **E’-H’**) flies, stained for DAPI (blue), anti-BAM (red), anti-CAL1 (green) and anti-VASA (not shown). Germ cells belonging to the 16-cell cyst chamber were selected based on the VASA marker and the lack of BAM signal in the control (**A’-D’**) and in the CAL1 RNAi (**E’-H’**). White Dotted line (**D** and **H**) indicates the end of the BAM positive region. At this stage, centromeres (CAL1 foci) cluster at the nucleolus. Star indicates the position of the terminal filament; 3 days old female flies; scale bar: 10 μm.

