## Supplemental figures for "CENP-A drives asymmetric cell division and maintains stem identity"

**Supplemental Information**

**Figure S1, related to Figure 2**

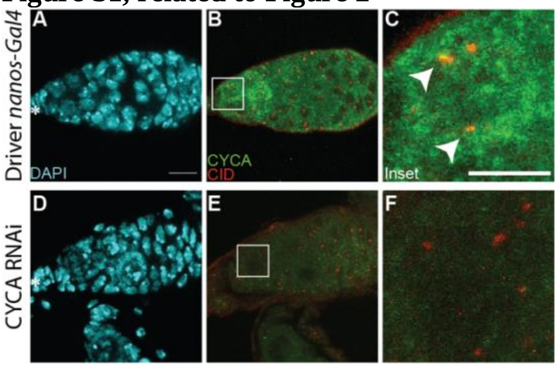

**Figure S2, related to Figure 3**

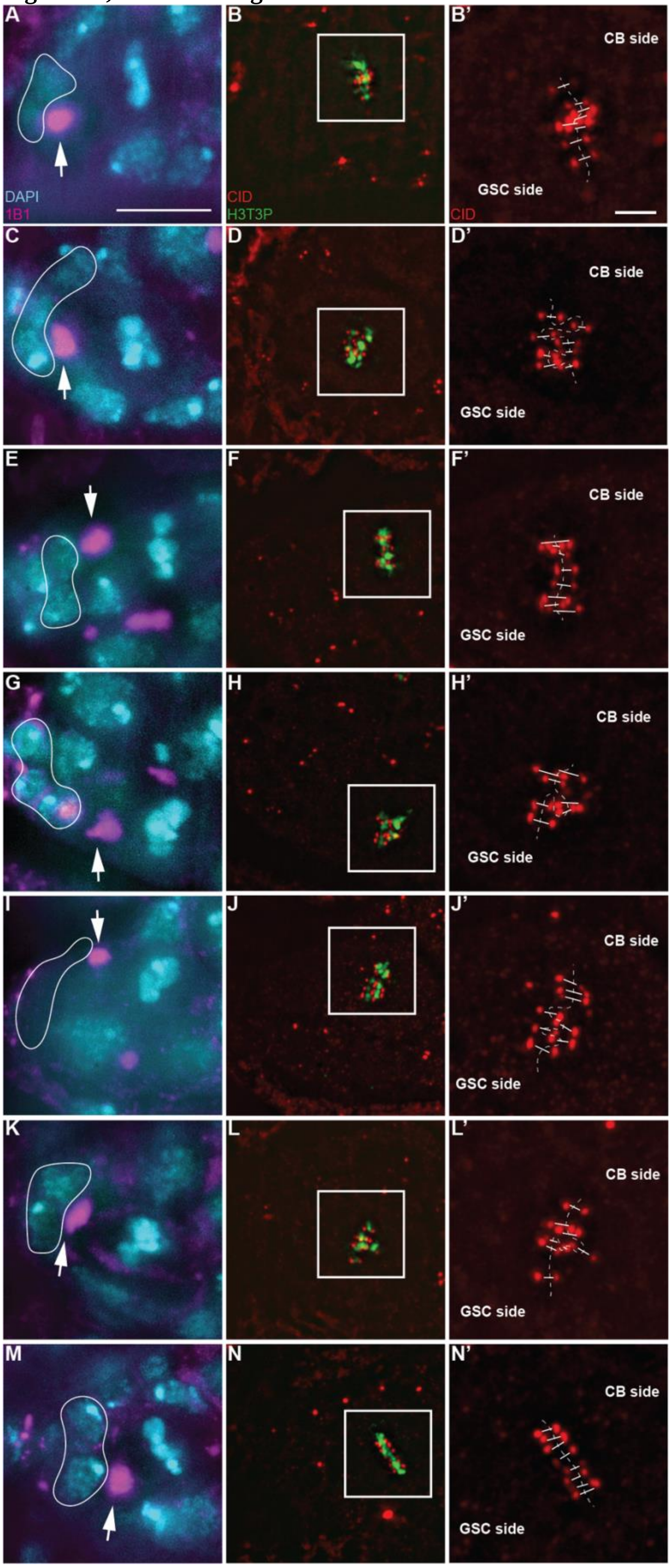

**Figure S3, related to Figure 4**

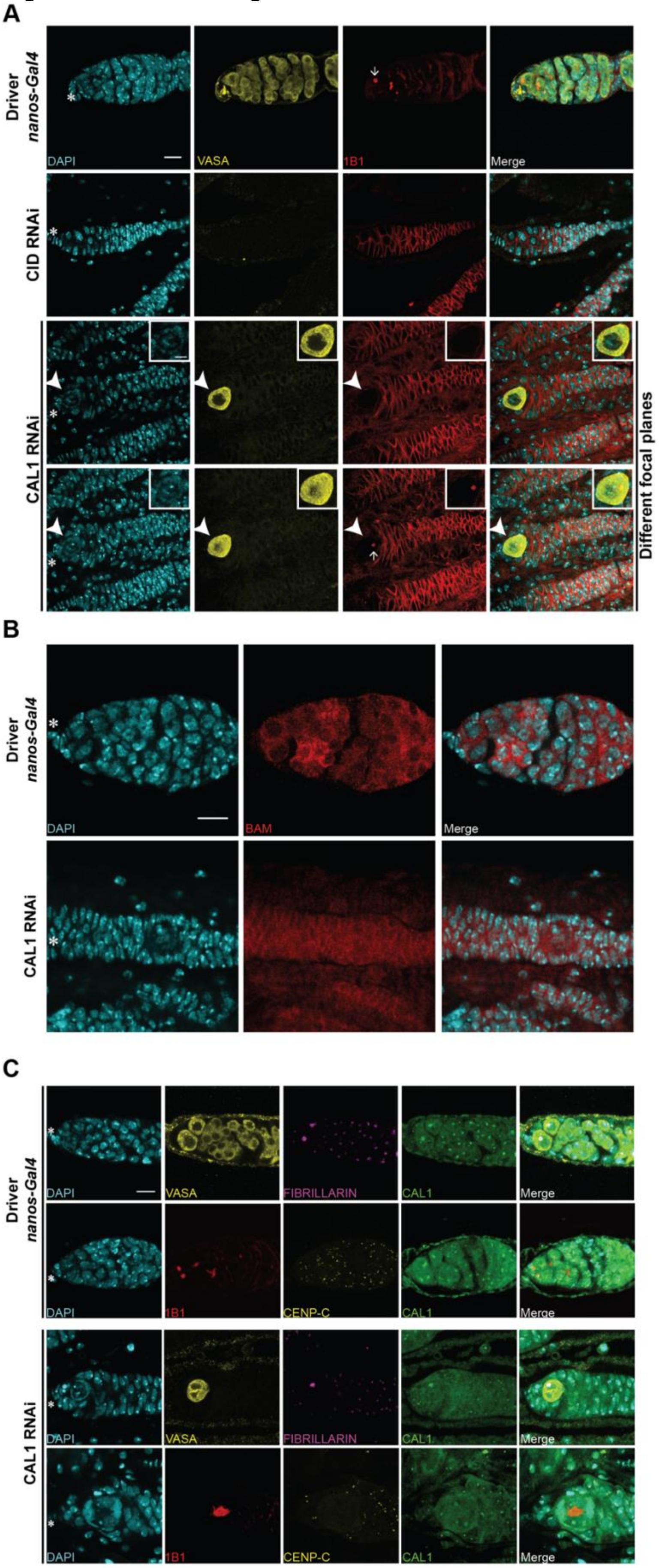

**Figure S4, related to Figure 5**

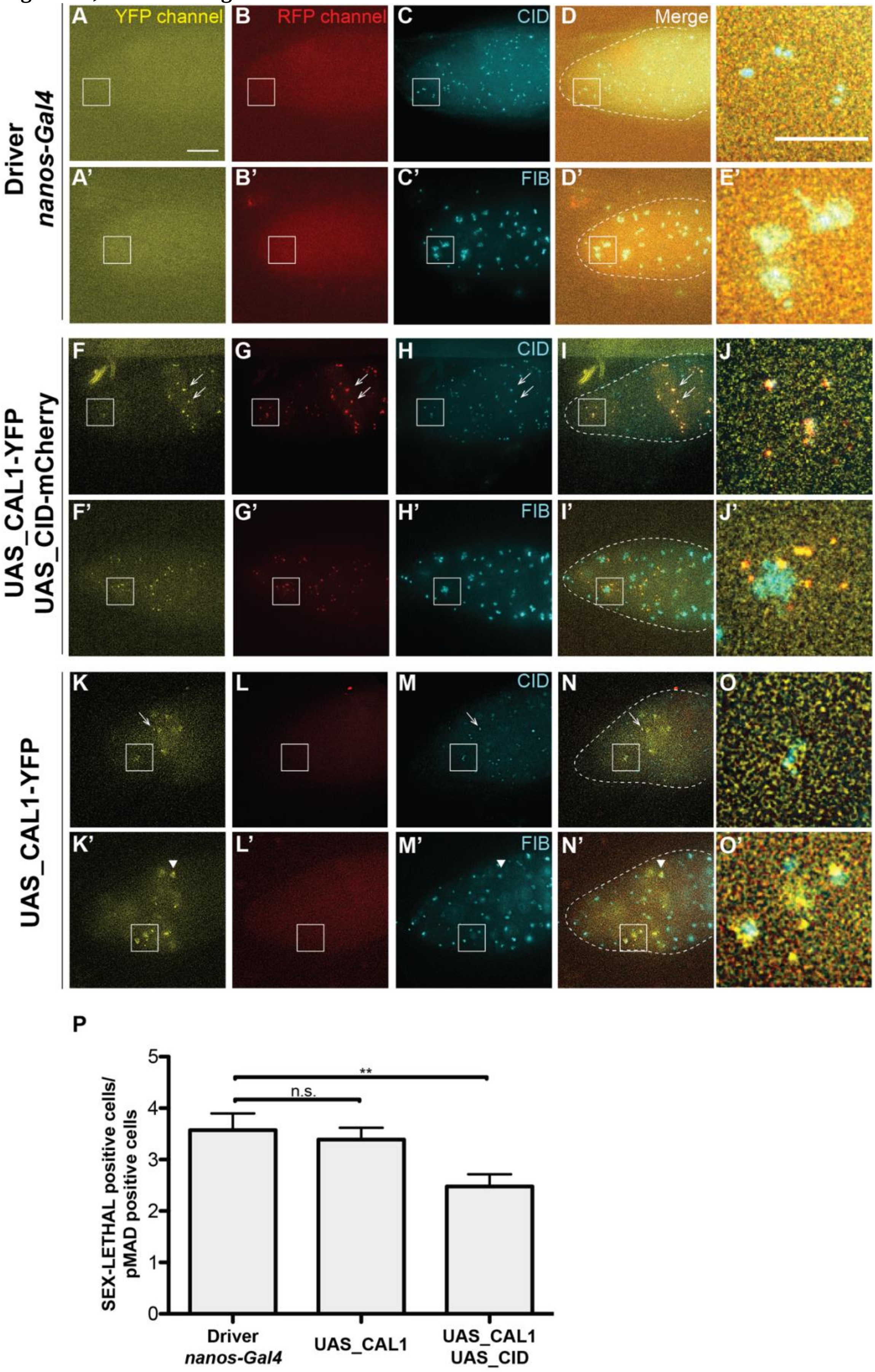

**Figure S5, related to Figure 6**

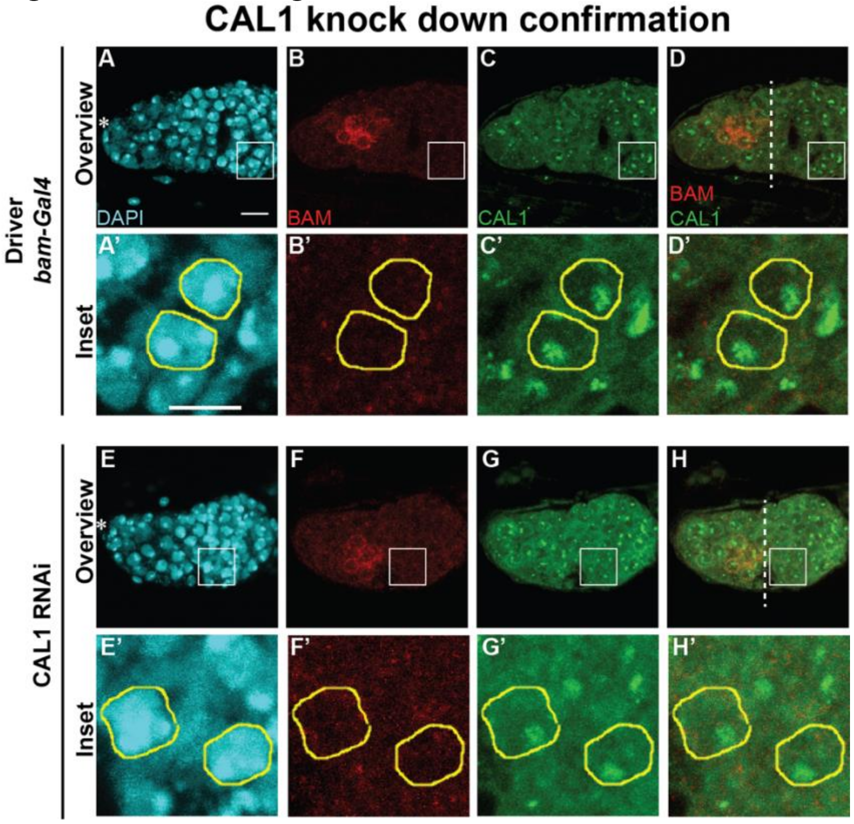
